# A locked immunometabolic switch underlies TREM2 R47H loss of function in human iPSC--derived microglia

**DOI:** 10.1101/766089

**Authors:** Thomas M Piers, Katharina Cosker, Anna Mallach, Gabriel Thomas Johnson, Rita Guerreiro, John Hardy, Jennifer M Pocock

## Abstract

Loss-of-function genetic variants of *triggering receptor expressed on myeloid cells 2 (TREM2)* are linked with an enhanced risk of developing dementias. Microglia, the resident immune cell of the brain, express TREM2 and microglial responses are implicated in dementia pathways. In a normal surveillance state, microglia use oxidative phosphorylation for their energy supply, but rely on the ability to undergo a metabolic switch to glycolysis to allow them to perform rapid plastic responses. We investigated the role of TREM2 on microglial metabolic function in human patient iPSC-derived-microglia expressing loss of function variants in TREM2. We show that these TREM2 variant iPSC-microglia, including the Alzheimer’s disease R47H risk variant, exhibit significant metabolic deficits including a reduced mitochondrial respiratory capacity and an inability to perform a glycolytic immunometabolic switch. We determined that dysregulated PPARγ/p38MAPK signalling underlies the observed phenotypic deficits in TREM2 variants and that activation of these pathways can ameliorate the metabolic deficit in these cells and consequently rescue critical microglial cellular function such as β-Amyloid phagocytosis. These findings have ramifications for microglial focussed-treatments in AD.

## Introduction

Microglia are the tissue resident macrophages and innate immune cells of the brain and need to rapidly respond to changes in their environment. Quiescent, surveillant innate immune cells such as microglia present in normal brain respond to activating stimuli by reprogramming metabolism, specifically, switching to favour glycolysis over oxidative phosphorylation (Kelly & O’Neill, 2015). Whilst glycolysis is an inefficient way to generate ATP, this switch enables microglia to undergo the rapid changes associated with their enhanced plasticity of function (Orihuela *et al*, 2016). It is also the preferential mechanism in proliferative cells to facilitate the uptake and incorporation of essential nutrients vital for the production of new cellular biomass (Vander Heiden *et al*, 2009).

In neurodegenerative diseases, aberrant microglial function has been increasingly recognised to contribute to disease progression however the influence of the cellular metabolic phenotype is not well understood. Missense mutations of *triggering receptor expressed on myeloid cells 2 (TREM2)* are associated with an enhanced risk of developing dementias including late onset Alzheimer’s disease (LOAD) (Guerreiro *et al*, 2013; Jonsson *et al*, 2013). In the CNS, TREM2 is exclusively expressed in microglia, and numerous studies have linked the disease-associated mutations to deficits in microglial function, including ligand binding/sensing, phagocytosis, and inflammatory responses (Kleinberger *et al*, 2014; Wang *et al*, 2015).

Much of the current work to elucidate the loss of function consequences of TREM2 variants in AD has employed the use of KO animal models and whilst a role for TREM2 has been described in microglial metabolism (Ulland et al 2017), it is not known whether disease-relevant variants also harbour metabolic deficits or the nature of any observed deficits. Here, we used human iPSC-derived microglia (iPS-Mg) generated from donors harbouring specific TREM2 mutations previously characterised as hypomorphic variants in Alzheimer’s disease and Nasu Hakola disease (NHD), and identified deficits in microglial metabolic regulation and associated functions. Furthermore, we identified for the first time that TREM2 variants are unable to carry out an immunometabolic switch to induce glycolysis and that this depends on PPARγ-p38MAPK-PFKFB3 signalling.

## Materials and Methods

### iPSC generation

Ethical permission for this study was obtained from the National Hospital for Neurology and Neurosurgery and the Institute of Neurology joint research ethics committee (study reference 09/H0716/64) or by the Ethics Committee of Istanbul Faculty of Medicine, Istanbul University (for collection of T66M mutant fibroblasts to Dr Ebba Lohmann). R47H heterozygous fibroblasts were acquired with a material transfer agreement between University College London and University of California Irvine Alzheimer’s Disease Research Center (UCI ADRC; M Blurton-Jones). Fibroblast reprogramming was performed by episomal plasmid nucleofection (Lonza) as previously described (Okita *et al*, 2011), using plasmids obtained from Addgene (#27077, #27078 and #27080). Nucleofected cultures were transferred to Essential 8 medium (Life Technologies) after 7 days *in vitro* (DIV) and individual colonies were picked after 25-30 DIV and CNV analysis performed. All iPSCs were maintained and routinely passaged in Essential 8 medium. Karyotype analysis was performed by The Doctors Laboratory (London, UK) (Supplementary Fig 2B-D). The R47H^hom^ line was a gene-edited isogenic of BIONi010-C, purchased from EBiSC (BIONi010-C7). Control iPSC lines used in this study are as follows: CTRL1 (kindly provided by Dr Selina Wray, UCL Institute of Neurology); CTRL2 (SBAD03, Stembancc); CTRL3 (SFC840, Stembancc); CTRL4 (BIONi010-C, EBiSC).

### iPSC-derived microglia (iPS-Mg)

Using our previously described protocol, iPSC-microglia (iPS-Mg) were generated (Xiang *et al*, 2018).

### Cellular Stress Proteome array

Cells were treated for 8 hours with 2-deoxyglucose (2-DG; 3mM) and cell lysates were prepared as per the manufacturer’s instructions (Proteome Profiler™ Human cell stress array; Bio-Techne). Total protein quantification was performed on aliquots of each treatment group for data normalisation purposes. Lysates were pooled from 3 independent experiments, according to *TREM2* genotype after basal or 2-DG treatment. Data were analysed using the Protein Array Analyser Palette plugin for ImageJ (Carpentier & Henault, 2010), and plotted as relative protein expression, normalised to total cellular protein levels.

### Immunoblotting

iPS-Mg were lysed in RIPA buffer (50 mM Tris, 150mM NaCl, 1% SDS, and 1% Triton X-100) containing 1x Halt™ protease and phosphatase inhibitor cocktail. Lysates were separated into soluble and insoluble (nuclear) fractions. Samples were resolved and transferred onto nitrocellulose membranes and incubated with primary and secondary antibodies (Table 1). Blotting was visualised on an Odyssey detection system (LiCor) and quantified using ImageJ software (www.imagej.nih.gov/ij).

**Table 1.**
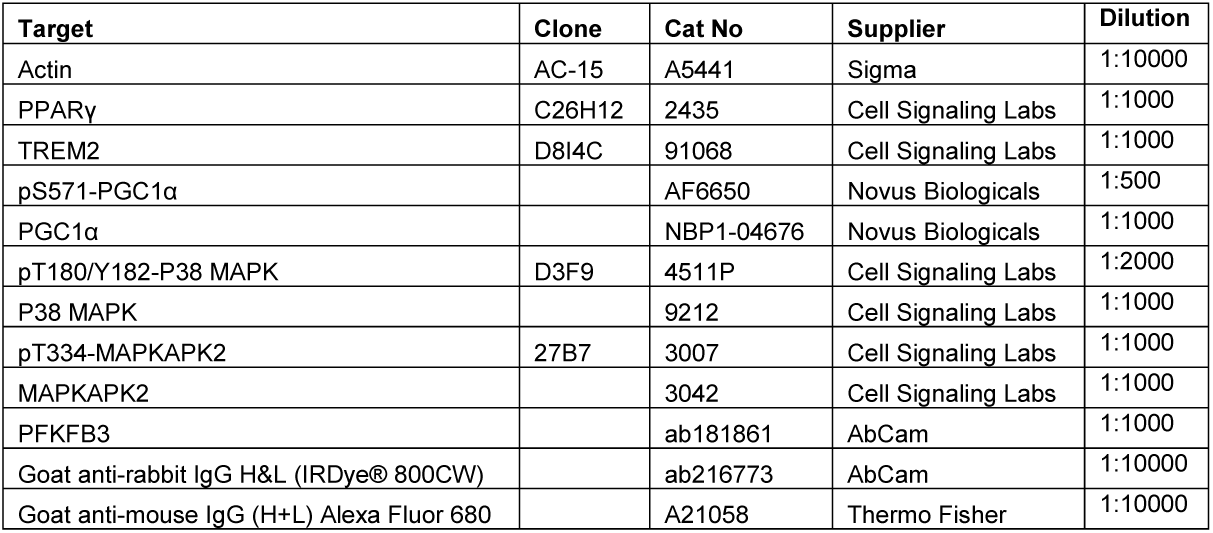
Primary and secondary antibody details.

| Target | Clone | Cat No | Supplier | Dilution |
| --- | --- | --- | --- | --- |
| Actin | AC-15 | A5441 | Sigma | 1:10000 |
| PPARγ | C26H12 | 2435 | Cell Signaling Labs | 1:1000 |
| TREM2 | D8I4C | 91068 | Cell Signaling Labs | 1:1000 |
| pS571-PGC1α | | AF6650 | Novus Biologicals | 1:500 |
| PGC1α | | NBP1-04676 | Novus Biologicals | 1:1000 |
| pT180/Y182-P38 MAPK | D3F9 | 4511P | Cell Signaling Labs | 1:2000 |
| P38 MAPK |  | 9212 | Cell Signaling Labs | 1:1000 |
| pT334-MAPKAPK2 | 27B7 | 3007 | Cell Signaling Labs | 1:1000 |
| MAPKAPK2 |  | 3042 | Cell Signaling Labs | 1:1000 |
| PFKFB3 |  | ab181861 | AbCam | 1:1000 |
| Goat anti-rabbit IgG H&L (IRDye® 800CW) |  | ab216773 | AbCam | 1:10000 |
| Goat anti-mouse IgG (H+L) Alexa Fluor 680 |  | A21058 | Thermo Fisher | 1:10000 |

### Live cell morphological staining

iPS-Mg were matured on 13mm glass coverslips. Cells were treated with 100ng/ml LPS and 10U/ml human IFNγ for 24 hours prior to staining. Prior to visualisation, cells were incubated with 1 µM Calcein-AM for 15 minutes at 37°C. Images were captured on a Zeiss Axioskop 2 fluorescence microscope and image analysis was carried out with Axiovison 4.8 and ImageJ software.

### Mitochondrial superoxide analysis

Basal levels of mitochondrial superoxide were measured by loading cells with MitoSOX**™** red superoxide indicator (Thermo Fisher) and analysed by flow cytometry. Briefly, a 5µM working concentration of MitoSOX**™** red was prepared in warmed FACs buffer (PBS + 0.5% BSA). Cells (400,000) were incubated with MitoSOX**™** red for 10 minutes at 37°C, protected from light. Cells were washed twice and resuspended in FACs buffer followed by flow cytometry (FL2; FACs Calibur, Beckton Dickinson). Incubation with Rotenone (100nM) for 30 minutes was used a positive control. Data were analysed using Flowing Software v2.5.1 (University of Turku).

### Mitochondrial number determination

To assess gross mitochondrial number in the iPS-Mg lines, flow cytometry analysis using MitoTracker**™** Green (Thermo Fisher) was performed. Briefly, a 200nM working concentration of MitoTracker**™** Green was prepared in warmed FACs buffer (PBS + 0.5% BSA). Cells (400,000) were incubated with MitoTracker**™** Green for 30 minutes at 37°C, protected from light. Cells were washed twice and resuspended in FACs buffer followed by flow cytometry (FL1; FACs Calibur, Beckton Dickinson). Data were analysed using Flowing Software v2.5.1 (University of Turku).

### PPARγ transcriptional activity

The DNA binding activity of PPARγ was detected in iPS-Mg nuclear extracts as previously described (Jiang *et al*, 2014) from untreated and groups treated with 2-DG (3mM) for 8 hours, using a PPARγ transcription factor assay kit, as per the manufacturer’s instructions (AbCam).

### Cellular respiration analysis

For real-time analysis of oxygen consumption rates (OCR) and extracellular acidification rates (ECAR), iPS-Mg were plated and matured on Seahorse cell culture microplates and analysed using a Seahorse XFe96 Analyser (Agilent Technologies). Cells were incubated overnight with or without pioglitazone (100nM) or GW0742 (100nM). Mito stress kits were used to analyse mitochondrial respiration and Glycolytic stress kits were used to analyse cellular glycolysis. Data were analysed using Wave v2.4.0.6 software (Agilent Technologies).

### 6-phosphofructokinase (PFK) activity assay

Activity of the glycolysis enzyme PFK was assessed in iPS-Mg cellular lysates after treatment with pioglitazone (100nM) for 24 hours +/- pre-treatment with SB202190 (100nM; 1 hour prior), as per the manufacturer’s instructions (AbCam).

### Aβ_1-42_ (HiLyte488) phagocytosis

Cells were plated and matured at a density of 20,000/well in 24 well plates (2 wells pooled/treatment group). Cells were treated with pioglitazone (100nM) +/- pre-treatment (1 hour prior) with SB202190 (100nM) or AZ-PFKFB3 (50nM) for 24 hours prior to FACs analysis. On the day of the experiment, cells were removed from the incubator to equilibrate to RT for 30 minutes and cytochalasin-D (CytoD; 100µM) was added to negative control groups. Aβ_1-42_ (HiLyte488; 100nM) was added to all groups except unstained groups and allowed to bind to phagocytic receptors for 30 minutes at RT. Grouped wells were then pooled into 2ml tubes, centrifuged at 300*g* for 3 minutes at RT, media aspirated, and the cell pellet resuspended in fresh RT maturation medium. Tubes were incubated at 37°C + 5% CO_2_ for 1 hour to initiate phagocytosis of bound Aβ_1-42_. Tubes were centrifuged at 300*g* for 3 minutes at RT, media was aspirated, and cell pellets were resuspended in PBS for FACs analysis (FL1; FACs Calibur, Beckton Dickinson). Data were analysed using FCS Express 6 Plus (DeNovo Software).

## Results

### TREM2 variant human iPS-Mg exhibit reduced oxidative phosphorylation and glycolytic capability

Using our previously described techniques (Xiang *et al*, 2018) we generated human iPS-Mg harbouring polymorphisms implicated in Alzheimer’s disease (AD; R47H) and in Nasu Hakola disease (NHD; T66M/W50C) that give rise to differential TREM2 protein glycosylation and cleavage (**Fig 1A**). Cellular respiration by Seahorse analysis of the oxygen consumption rate (*OCR*) and extracellular acidification rate (*ECAR*) was investigated in all available lines. Under mitochondrial stress and glycolytic stress, deficits were observed in oxidative phosphorylation (**Fig 1Bi**) and glycolytic function (**Fig 1Bii**) respectively. Specifically, levels of maximal respiration (**Fig 1C**), glycolysis (**Fig 1D**) and glycolytic capacity (**Fig 1E**) were significantly reduced in all iPS-Mg harbouring TREM2 variants compared with control lines. We confirmed that respiratory deficits were not due to underlying differences in the parental iPSC lines (**Fig 1F**); the results indicate that all lines prior to differentiation (and before TREM2 expression) displayed metabolic phenotypes independent of TREM2 genotype. We also measured mitochondrial number, since it has been reported that TREM^-/-^ mice exhibit reduced mitochondrial mass (Ulland et al., 2017), however we found no difference as measured by MitoTracker Green between control and TREM2 variant iPS-Mg lines (**Fig 1G**). Instead, production of mitochondrial superoxide assessed by MitoSOX red was increased in both AD R47H and NHD TREM2 iPS-Mg (**Fig 1Hi & Hii**) suggesting dysregulation of basal mitochondrial function in TREM2 mutant carriers.

**Figure 1.**
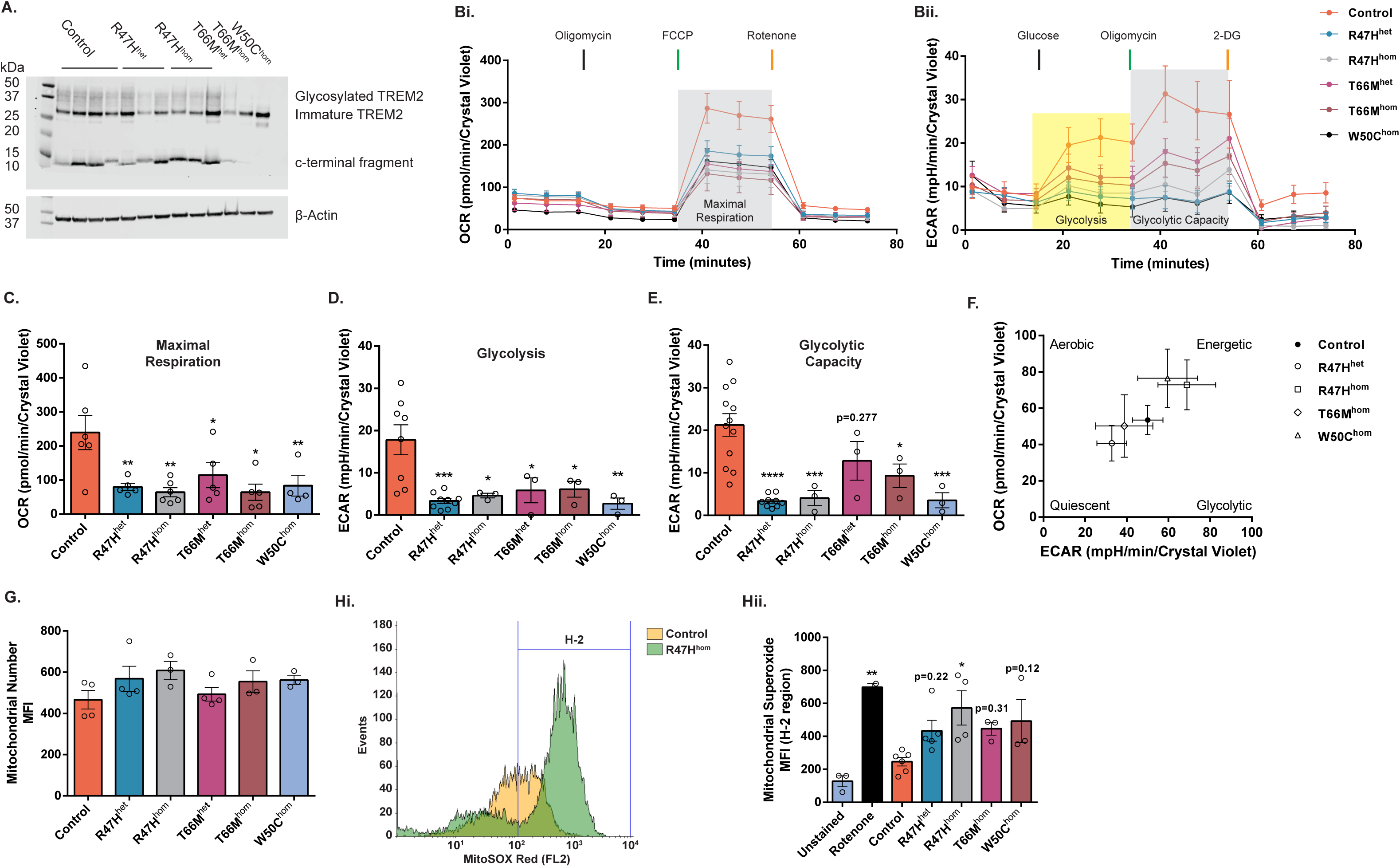
Significant deficits in cellular respiration are observed in iPS-Mg from TREM2 hypomorphic lines. **A**. Representative western blotting of TREM2 protein levels in control and variant TREM2 lines, confirming previously published differences in TREM2 glycosylation and cleavage patterns. **B**. The oxygen consumption rate (OCR) (**Bi**) and the extracellular acidification rate (ECAR) (**Bii**) was analysed in control iPS-Mg and TREM2 mutant iPS-Mg to assay mitochondrial oxidative function and the glycolytic function of the lines, respectively. **C**. Analysis of iPS-Mg maximal respiration, post FCCP injection, identified significant deficits in the OCR of TREM2 hypomorphic lines, when compared with controls. **D**. Analysis of iPS-Mg glycolysis, post glucose injection, identified significant deficits in the ECAR of TREM2 hypomorphic lines, when compared with controls. **E**. Analysis of iPS-Mg glycolytic capacity, post oligomycin injection, identified significant deficits in the ECAR of TREM2 hypomorphic lines, when compared with controls. **F**. The basal metabolic phenotype of iPSC parental lines used to develop iPS-Mg are independent of TREM2 genotype. **G**. MitoTracker™ Green flow cytometry of control and TREM2 hypomorphic iPS-Mg suggested the observed deficits in cellular respiration were not due to differences in mitochondrial number. Plotted as mean fluorescent intensity (MFI). **H**. Representative histogram of MitoSOX™ red flow cytometry in control and R47H^hom^ iPS-Mg (**Hi**) and quantification of basal mitochondrial reactive oxygen species (mean fluorescence intensity, MFI ± SEM) (**Hii**) show significantly higher levels in TREM2 hypomorphic mutants when compared to control levels Data information: In (**B-G, Hiii**) data are represented as mean ± SEM (*n*≥3). Statistical significance was addressed using 1-way ANOVA with Bonferroni’s multiple comparison test to compare control iPS-Mg with TREM2 hypomorphic iPS-Mg lines or positive control, \**P*<0.05; \*\**P*<0.01; \*\*\**P*<0.005; \*\*\*\**P*<0.001.

### iPS-Mg with TREM2 hypomorphic variants fail to “switch” to glycolysis following an inflammatory challenge

Given the energetic deficits in TREM2 variant iPS-Mg in response to mitochondrial and glycolytic stress, we investigated how these cells respond to an inflammatory challenge. We found that following exposure to lipopolysaccaride (LPS) and interferon-gamma (IFNγ), control iPS-Mg undergo morphological changes, specifically increased cellular elongation and a reduction in ramified processes (**Fig 2A)**, and show substantial release of the pro-inflammatory cytokine TNFα (**Fig 2B**). In TREM2 variant iPS-Mg, the morphological changes are less dramatic (**Fig 2A**) and they exhibit a significant reduction in TNFα release compared with control lines (**Fig 2B**), indicating a deficit in the microglial response to the inflammatory stimulus. Microglia respond to pro-inflammatory stimuli by initiating a metabolic switch from oxidative phosphorylation to glycolysis. Seahorse analysis was again employed to measure *OCR* and *ECAR*, this time following exposure to LPS and IFNγ. Control iPS-Mg showed a robust shift from low quiescent to high energetic respiration in response to LPS and IFNγ (**Fig 2C**). In contrast, TREM2 variant iPS-Mg from AD and NHD lines remained “stuck” in a quiescent respiratory state (**Fig 2C**). When glycolysis was blocked with 2-deoxyglucose (2-DG, 3mM) prior to LPS/IFNγ, control iPS-Mg are still able to generate aerobic energy (**Fig 2D**) to compensate. Interestingly, R47H^het^ iPS-Mg were also able to induce aerobic respiration following inhibition of glycolysis, unlike the other TREM2 homozygous variants that remained in a quiescent state (**Fig 2D**).

**Figure 2.**
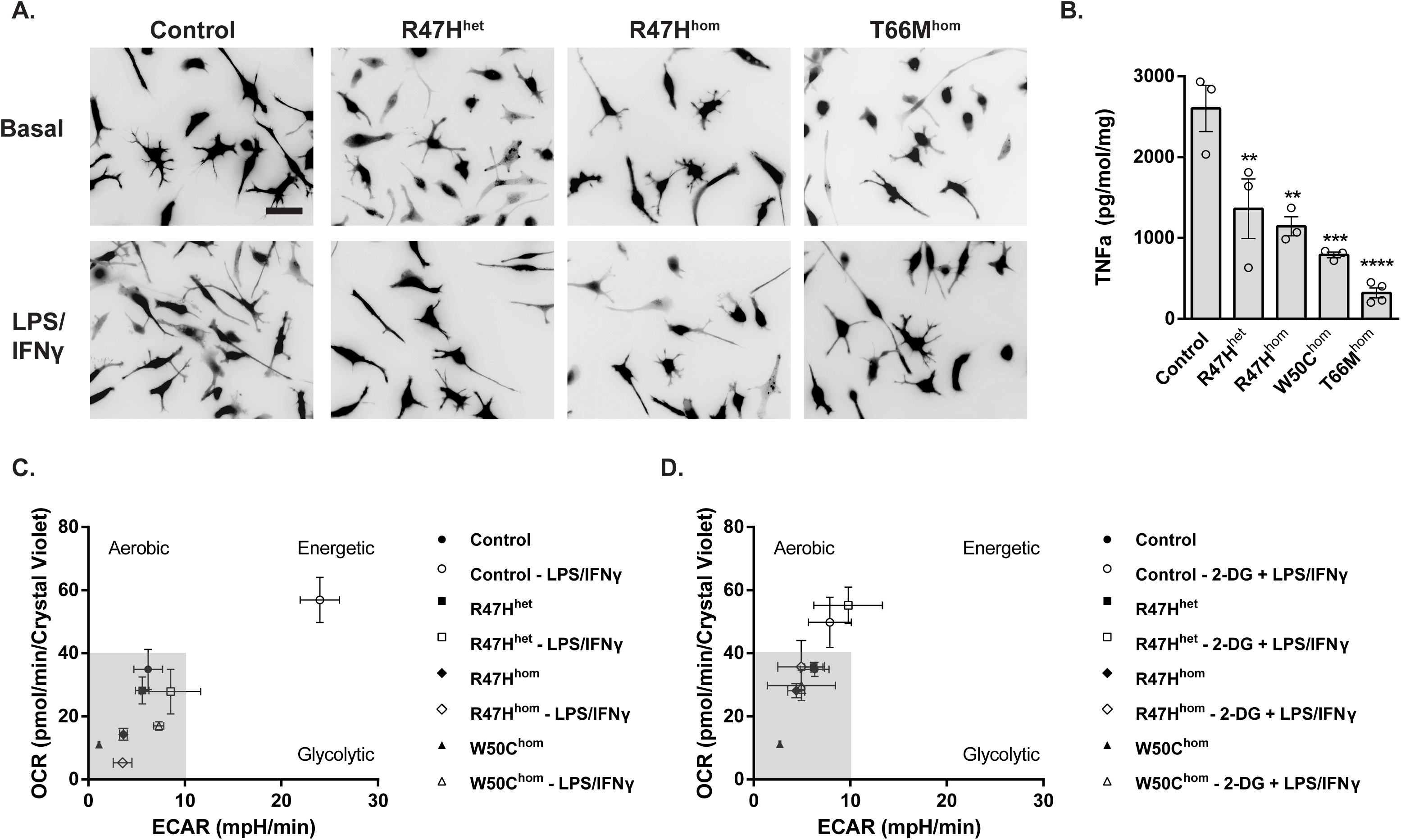
Activated morphologies and release of TNFα were reduced in TREM2 hypomorphic iPS-Mg. **A**. Representative images of iPS-Mg after LPS/IFNγ treatment shows a reduction of activated morphologies in TREM2 mutant lines **B**. ELISA analysis of LPS/IFNγ treated iPS-Mg identified a significant reduction in TNFα release from TREM2 mutant lines **C**. Metabolic phenotypes of iPS-Mg after LPS/IFNγ treatment show a loss of a metabolic switch in TREM2 mutant lines **D**. Metabolic phenotypes of iPS-Mg after inhibition of glycolysis and LPS/IFNγ treatment show a loss of switch to active oxidative phosphorylation in TREM2 homozygous mutant lines Data information: In (**A**), scale bar: 50 μm. In (**B-D**), data are presented as mean ±SEM (*n*≥3). Statistical significance was addressed using 1-way ANOVA with Bonferroni’s multiple comparison test to compare control iPS-Mg with TREM2 hypomorphic iPS-Mg lines, \*\**P*<0.01; \*\*\**P*<0.005; \*\*\*\**P*<0.001.

### TREM2 variant iPS-Mg induce cellular stress pathways in response to glycolytic inhibition and exhibit reduced PPARγ signalling

In order to investigate the comparable aerobic responses after glycolytic inhibition, but underlying phenotypic differences between the control and patient AD R47H^het^ iPS-Mg, cells were treated with 2- DG and probed for changes in key cell stress pathways. In 2-DG-treated R47H^het^ iPS-Mg we identified enhanced levels of cell stress proteins that are linked to energy metabolism and/or mitochondrial function, including HIF1α, cytochrome-c, Bcl-2, Cited-2, NF-κB and p38α MAPK (**Fig 3Ai & 3Aii**). PGC- 1α, the master regulator of mitochondrial biogenesis and energy metabolism (Liang & Ward, 2006; Austin & St-Pierre, 2012), is downstream of a number of these pathways and so we examined phosphorylation levels at serine 571, which negatively regulates PGC-1α activity in control and TREM2 variant iPS-Mg. We found that phosphorylation at serine 571, was significantly enhanced in TREM2 variant iPS-Mg, suggesting that PGC-1α activity is downregulated in TREM2 variant cells (**Fig 3Bi & Bii**). PGC-1α is also a co-activator with the nuclear receptor and transcription factor PPARγ widely expressed in inflammatory cells such as microglia and macrophages as well as adipose tissue where it controls inflammation, lipid metabolism and glucose homeostasis (Corona & Duchen, 2016). Following 2-DG treatment, we found increased PPARγ transcriptional activity in control iPS-Mg but no increase in AD R47H^het^ or NHD T66M^hom^ iPS-Mg (**Fig 3C**) despite the increase in cell stress pathways in AD R47H^het^ iPS-Mg. When we looked at total PPARγ protein levels in iPS-Mg, we found that PPARγ protein levels were significantly reduced in TREM2 hypomorphs, either at basal (T66M^hom^/W50C^hom^) or after glycolytic inhibition with 2-DG (R47H^het/hom^ and T66M^het^) (**Fig 3Di & Dii**).

**Figure 3.**
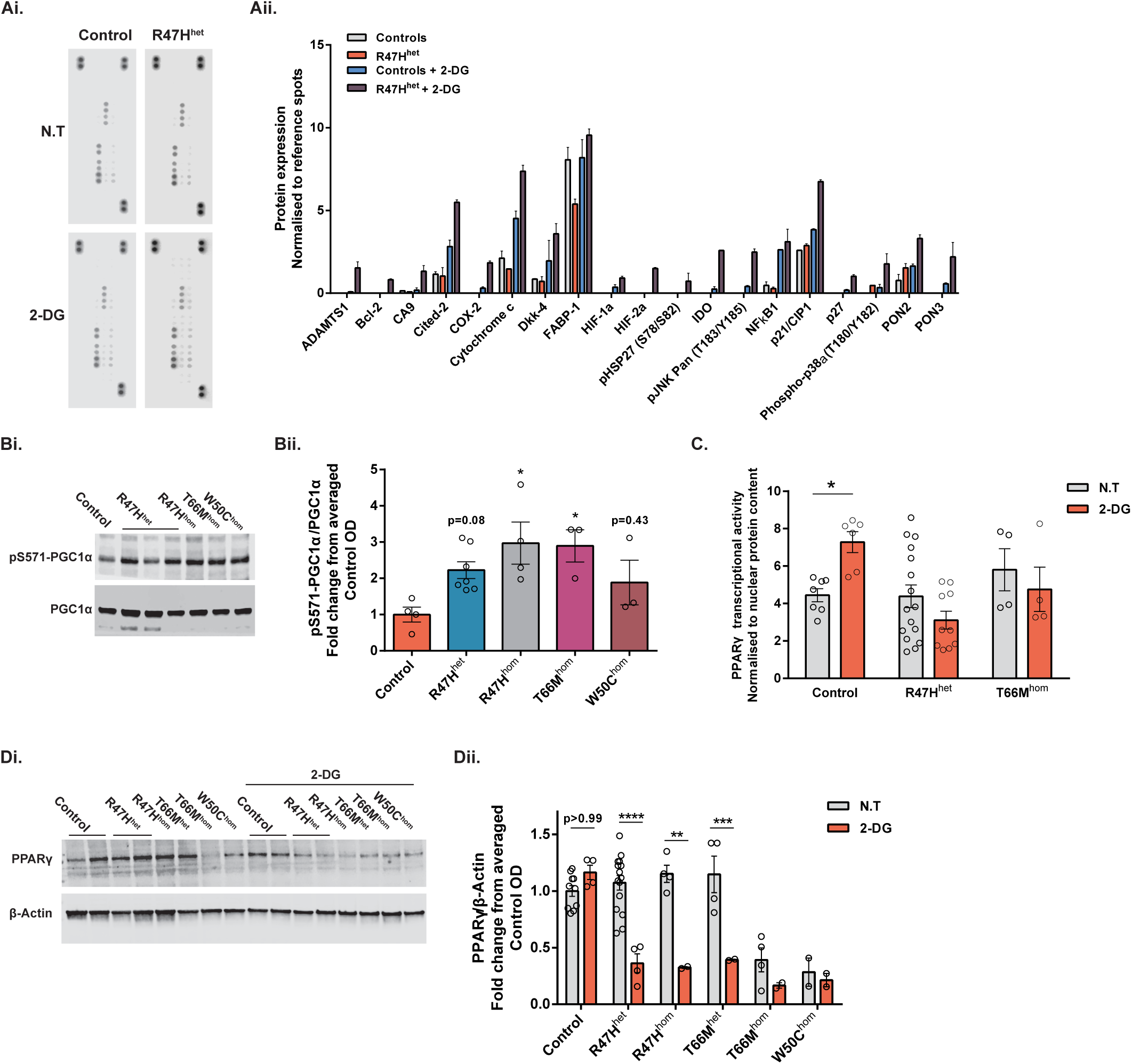
iPS-Mg with TREM2 hypomorphs have enhanced cellular stress indicators and aberrant PPARγ signalling. **A**. Representative cellular stress proteome array dot blots from control and R47H^het^ iPS-Mg cell lysates +/- 2-DG (3mM) treatment (**Ai**) and quantification of cell stress proteome array show increased levels of a number of cellular and mitochondrial associated stress proteins in R47H^het^ iPS-Mg lysates after 2- DG treatment (**Aii**). **B**. Representative western blotting of phospho-S571-PGC1α and total PGC1α protein levels in control and mutant TREM2 iPS-Mg (**Bi**) and quantification of the protein levels identified enhanced levels of phospho-S571-PGC1α in TREM2 mutant expressing iPS-Mg lines (**Bii**). **C**. Cellular stress induced by 2-DG (3mM) enhanced PPARγ transcriptional activity in control iPS-Mg but not in R47H^het^ or T66M^hom^ iPS-Mg; control: p=0.0372, R47H^het^: p=0.3352, T66M^hom^: p>0.9999; n=4- 16, normalised to nuclear protein content ± SEM. **D**. Representative western blotting of PPARγ and β-Actin protein levels in control and mutant TREM2 iPS-Mg after 2-DG treatment (**Di**) and quantification identified reductions in PPARγ protein levels in TREM2 mutant iPS-Mg after 2-DG treatment, or in the case of T66Mhom and W50Chom, reduced basal levels of the protein compared to control levels (**Dii**). Data information: In (**Aii, Bii, C, Dii**), data are presented as mean ± SEM (*n*≥3). Statistical significance was addressed using 1-way ANOVA with Bonferroni’s multiple comparison test to compare control iPS-Mg with TREM2 hypomorphic iPS-Mg lines (**Bii**) or 2-way ANOVA with Bonferroni’s multiple comparison test to compare non-treated and 2-DG groups (**C, Dii**), \**P*<0.05; \*\**P*<0.01; \*\*\**P*<0.005; \*\*\*\**P*<0.001.

### A PPARγ agonist can rescue energy deficits and the metabolic glycolytic switch in TREM2 variant iPS-Mg

We set out to determine whether dysregulation of PPARγ signalling was the cause of the inability of TREM2 hypomorphs to switch to an energetic phenotype and whether this could be rescued by targeting PPARγ activation. We thus investigated whether oxidative phosphorylation and glycolysis could be modulated through PPARγ agonism; pre-incubation with the PPARγ agonist pioglitazone significantly attenuated the observed energy deficits in maximal respiration **(Fig 4Ai & 4Aii**), glycolysis (**Fig 4Bi & 4Bii**), and glycolytic capacity (**Fig 4Bi & 4Biii**) in iPS-Mg from R47H^het/hom^ carriers, but interestingly did not significantly enhance maximal metabolic respiration in control iPS-Mg, suggesting that metabolic function in these cells was already optimal or saturated. In the case of the T66M^hom^, or W50C^hom^ TREM2 hypomorphic iPS-Mg, whilst there was a trend to increased respiration with pioglitazone, this was not significant (**Fig 4Aii**) and pioglitazone did not enhance glycolysis in these cells either (**Fig 4Bii & 4iii**). Since the R47H variant expressing iPS-Mg were most amenable to rescue with pioglitazone treatment, we examined whether this activation could promote the switch to an energetic phenotype following inflammatory stimulation with LPS and IFNγ; indeed this was found to be the case (**Fig 4C**) suggesting we can reverse the metabolic deficit in these cells.

**Figure 4.**
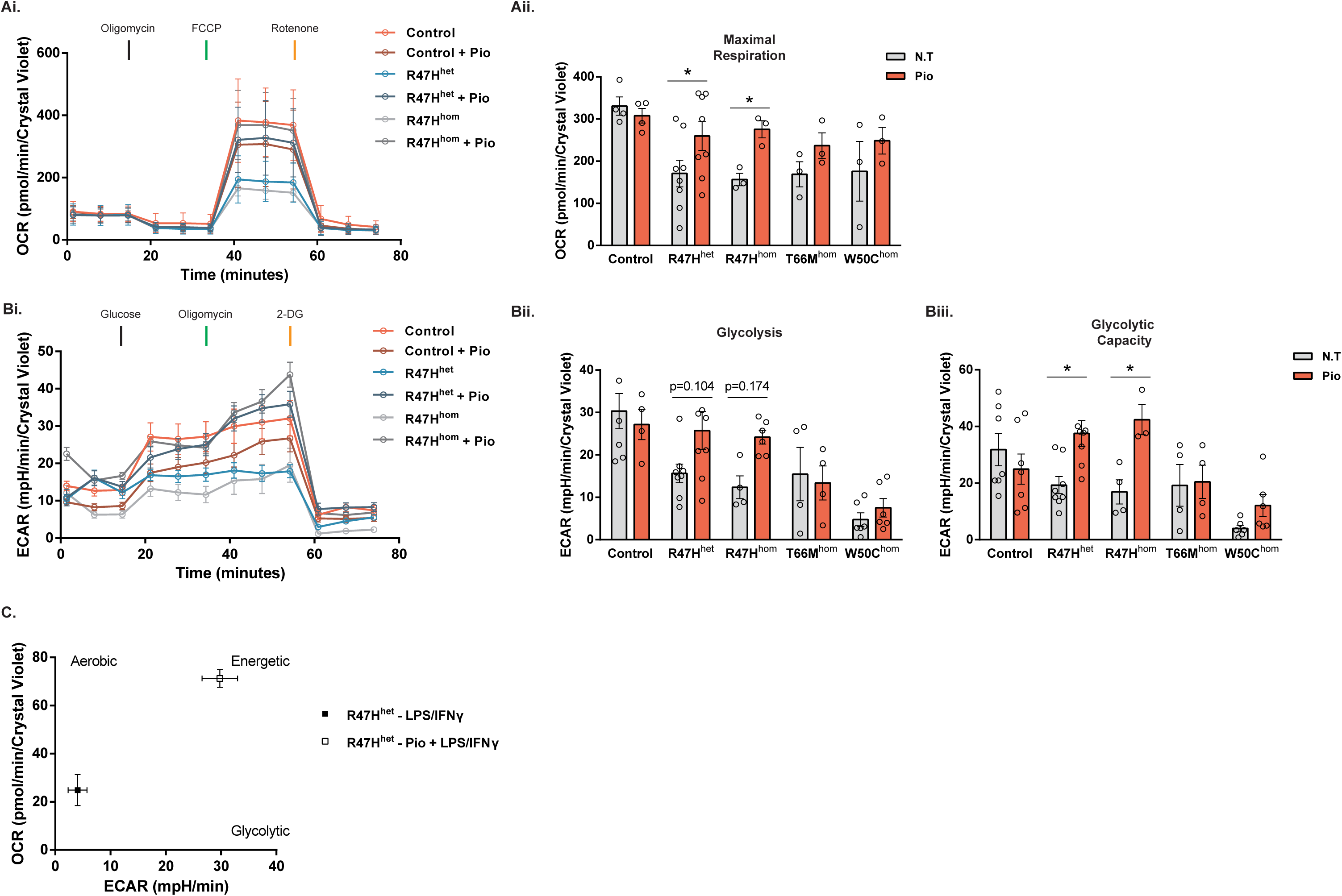
Activation of PPARγ signalling rescues cellular respiration and metabolic phenotype switch in AD-associated R47H iPS-Mg lines. **A**. Complete traces of the control and R47H hypomorphic iPS-Mg mitochondrial stress test pre-treated with 100nM pioglitazone (**Ai**). Pre-treatment with pioglitazone reversed the maximal respiration deficits observed in R47H iPS-Mg lines when compared with non-treated groups (**Aii**). **B**. Complete traces of the control and R47H hypomorphic iPS-Mg glycolytic stress test pre-treated with 100nM pioglitazone (**Bi**). Pre-treatment with pioglitazone partially reversed the glycolysis deficits (**Bii**) and significantly reversed glycolytic capacity deficits (**Biii**) in R47H lines. **C**. The LPS/IFNγ-induced metabolic switch is rescued in R47H^het^ iPS-Mg after activation of PPARγ. Data information: In (**A-C**), data are presented as mean ± SEM (*n*≥3). Statistical significance was addressed using 2-way ANOVA with Bonferroni’s multiple comparison test to compare non-treated with treated groups within genotypes, \**P*<0.05.

### Pioglitazone rescues the energy deficits in TREM2 variant iPS-Mg through p38-MAPK and PFKFB3 signalling

To further investigate the mechanisms by which pioglitazone can initiate the switch to glycolysis in TREM2 R47H variants, we probed the involvement of the PPARγ target PFKFB3 (Huo et al., 2010), a key regulatory enzyme in glycolysis (Rodríguez-García *et al*, 2017; Finucane *et al*, 2019). PFKFB3 regulates glucose metabolism via synthesis of fructose-2,6-bisphosphate, a potent allosteric activator of 6-phosphofructo-1-kinase (PFK-1), which catalyses the commitment step of glycolysis through conversion of fructose 6-phosphate and ATP to fructose 1,6-biphosphate and ADP (Li *et al*, 2018). The signaling pathway leading to activation of PFKFB3 is through p38-MAPK dependent phosphorylation of MK2 and subsequent phosphorylation of PFKFB3 for induction of the glycolytic switch (Bolaños, 2013; Novellasdemunt *et al*, 2013). We again used LPS/IFNγ to induce glycolysis in iPS-Mg and found increased phosphorylation of MK2 and total protein levels of PFK3B3 that is dependent on p38-MAPK activity. However, in R47H^hom^ iPS-Mg there is a substantial deficit in the ability of LPS/IFNγ to induce phospho-MK2, again suggesting these cells cannot induce glycolysis after an inflammatory challenge. Given that p38-MAPK was increased in R47H in response to glycolytic stress (Fig 3A), it was surprising that p38-MAPK-dependent phosphorylation of MK2 was decreased in response to LPS/IFNγ. We therefore investigated the effect of PPARγ activation by pioglitazone on basal p38MAPK in unstimulated iPS-Mg. We found strong induction of pT180/Y182- p38MAPK by western blot after pioglitazone exposure (100nM) in TREM2 variant iPS-Mg compared with control lines, suggesting that PPARγ induces a significant activation of the p38-MAPK pathway to rescue glycolysis in TREM2 lines that is not observed in control lines. Activation of PPARδ/β by GW0742 was ineffective at increasing p38MAPK phosphorylation (**Fig 5Bii**), suggesting this pathway is specific to PPARγ. We also probed PFK-1 activity downstream of PFKFB3 and show that pioglitazone is able to enhance PFK-1 activity, but only in the TREM2 hypomorphic lines and that this is p38-MAPK dependent (**Fig 5Ci-Ciii**), again supporting the idea that TREM2 variants can be rescued via activation of a p38-MAPK pathway to induce glycolysis. Indeed, when we probed glycolysis by Seahorse in R47H^hom^ iPS-Mg, inhibition of p38MAPK blocked the rescue of cellular glycolytic function by pioglitazone activation of PPARγ (**Fig 5D**).

**Figure 5.**
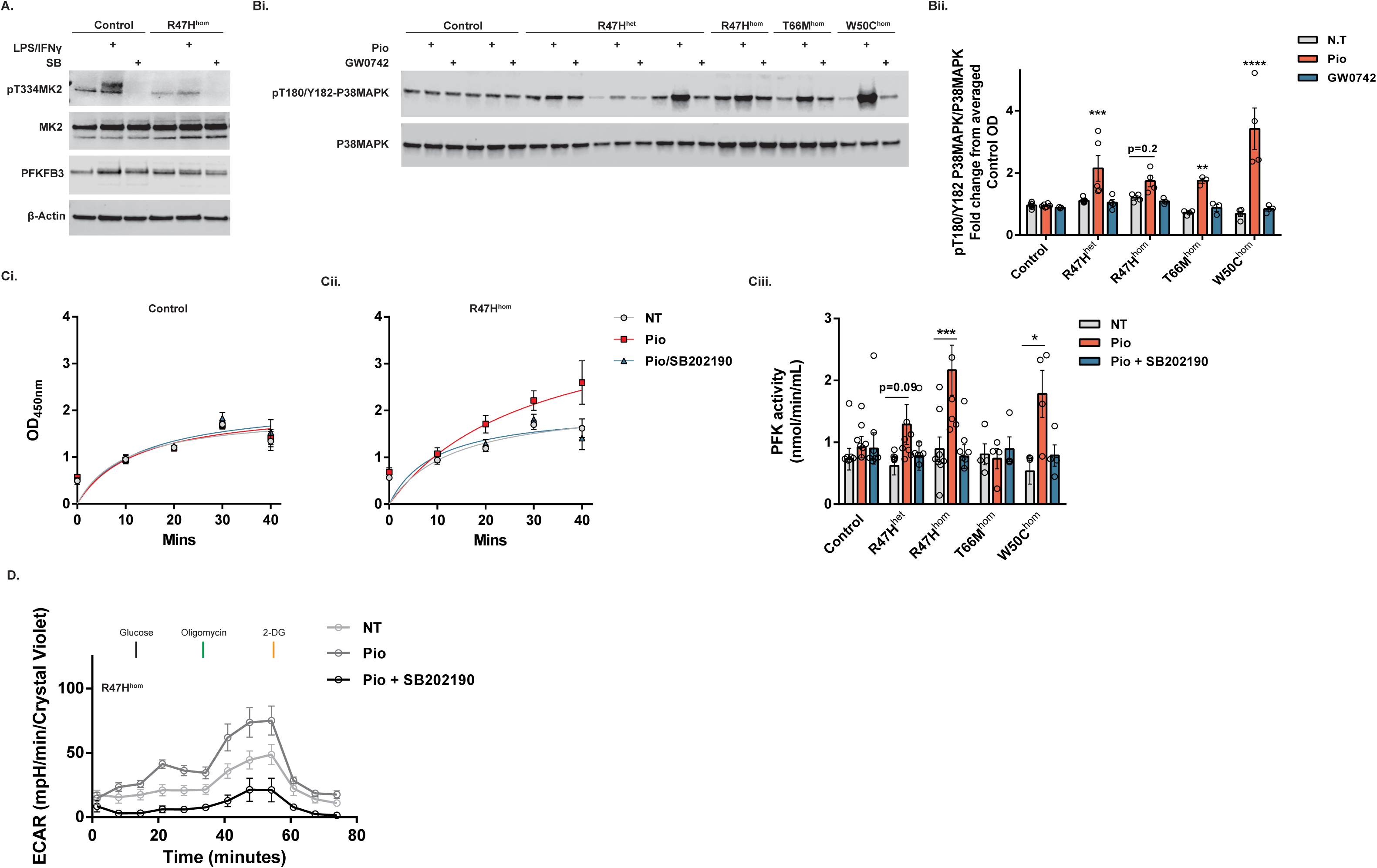
Pioglitazone rescues the energy deficits in TREM2 variant iPS-Mg through p38-MAPK and PFKFB3 signalling. **A**. The enhanced phosphorylation of MK2 and increase in PFKFB3 total protein in control iPS-Mg after pro-inflammatory induction is dependent on p38MAPK, and is reduced in TREM2 R47H iPS-Mg. **B**. Representative western blotting of phospho-T180/Y182-P38MAPK and total P38MAPK protein levels in control and mutant TREM2 iPS-Mg after pioglitazone or GW0742 treatment (**Bi**) and quantification identified enhanced levels of phospho-T180/Y182-P38MAPK after pioglitazone treatment specifically in TREM2 hypomorphic iPS-Mg lines (**Bii**). **C**. PFK1 enzyme activity is specifically enhanced by pioglitazone in TREM2 hypomorphic lines when compared to control iPS-Mg, and is dependent on p38MAPK (**Ci-Ciii**). **D**. Complete traces of the R47H^hom^ iPS-Mg glycolytic stress test show pioglitazone is able to enhance glycolysis and glycolytic function in a P38MAPK-dependent manner. Data information: In (**Bii-D**), data are presented as mean ± SEM (*n*≥3). Statistical significance was addressed using 2-way ANOVA with Bonferroni’s multiple comparison test to compare pioglitazone treatment of each TREM2 variant with the corresponding non-treated group, \**P*<0.05; \*\*\**P*<0.005; \*\*\*\**P*<0.001.

### Activation of the metabolic glycolytic switch by pioglitazone rescues the deficit in phagocytosis of Aβ_1-42_ in TREM2 variant iPS-Mg

Since pioglitazone can rescue the metabolic switch to glycolysis in TREM2 variant iPS-Mg, we asked whether PPARγ-p38MAPK/PFKFB3 activation can also rescue TREM2-dependent cellular functions. TREM2 has been shown to be important for phagocytosis, a crucial function of microglia that is found to be aberrant in neurodegenerative diseases such as Alzheimer’s (Kleinberger *et al*, 2014; Yeh *et al*, 2016). We have shown previously that NHD-associated TREM2 variant iPS-Mg have a deficit in phagocytosis of apoptotic cells (Garcia-Reitboeck *et al*, 2018) however this is not observed in R47H (data not shown). We therefore investigated the ability of TREM2 variants to clear amyloid beta (Aβ), which accumulates in plaques in AD and is the major pathological hallmark of the disease. Here we show that the R47H and the NHD variants exhibit a substantial deficit in their ability to phagocytose oligomeric Aβ_1-42_ compared with control cells (**Fig 6A-C**). Since phagocytosis has been shown to be driven in macrophages by glycolytic metabolism (Jiang *et al*, 2016), we determined whether pioglitazone could reverse the reduced phagocytosis of Aβ in TREM2 variant iPS-Mg. Activation of PPARγ by pioglitazone significantly rescues the TREM2 variant deficit in R47H lines, and this increase in phagocytosis is dependent on p38MAPK signalling (**Fig 6A & 6B**) and PFKFB3 activity (**Fig 6A & 6C**). These data suggest that the recovery of the glycolytic switch by activation of PPARγ-p38MAPK/PFKFB3 signalling is sufficient to rescue TREM2-dependent microglial functions such as phagocytosis of Aβ in AD.

**Figure 6.**
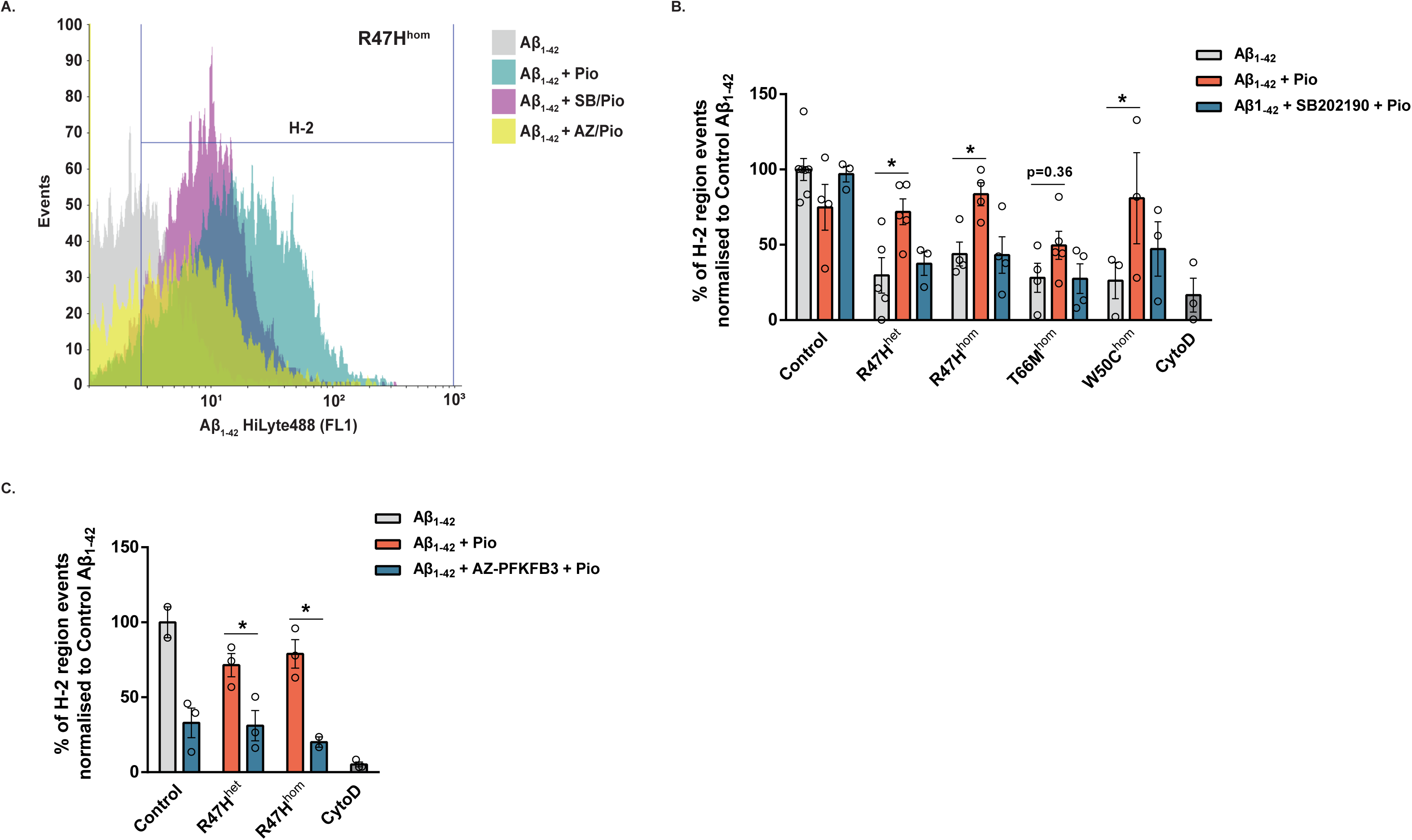
PPARγ mediated rescue of microglial functional deficits is dependent on p38MAPK signalling. **A**. Representative histogram of R47H^hom^ iPS-Mg-mediated phagocytosis of Aβ_1-42_ HiLyte488 after treatment with pioglitazone ± P38MAPK or PFKFB3 inhibition **B**. Quantification of Aβ_1-42_ phagocytosis in iPS-Mg identified that pioglitazone treatment enhanced phagocytosis in TREM2 variant lines and inhibition of P38MAPK prevented the observed rescue. **C**. Quantification of Aβ_1-42_ phagocytosis in control and R47H variant iPS-Mg identified that inhibition of PFKFB3 prevented the pioglitazone-mediated rescue of phagocytosis in the TREM2 variant lines. Data information: In (**B-C**), data are presented as mean ± SEM (*n*≥2). Statistical significance was addressed using 2-way ANOVA with Bonferroni’s multiple comparison test to compare Aβ_1-42_ + pioglitazone treatment of each TREM2 variant to the corresponding Aβ_1-42_ group (**B**), or to compare Aβ_1-42_ + pioglitazone treatment to Aβ_1-42_ + AZ-PFKFB3 inhibition + pioglitazone treatment in each corresponding group (**C**), \**P*<0.05.

## Discussion

Recent findings in mouse models suggest that TREM2 plays a role in maintaining microglial metabolic fitness (Ulland *et al*, 2017) and that TREM2-modulates the metabolic homeostasis of adipose tissue associated macrophages specifically the homeostasis of glucose, insulin, cholesterol, HDL and LDL (Jaitin *et al*, 2019) as well as disease-associated microglia in AD (Keren-Shaul *et al*, 2017).

Our data point to significant deficits in iPS-Mg harbouring TREM2 hypomorphic variants when in a basal homeostatic state. Specifically, these lines exhibit reduced maximal mitochondrial respiratory capacity as well as reduced glycolytic capacity when compared with common variant controls. We confirm these deficits correspond to the presence of TREM2 variants and are not an inherent property of the iPSC since the corresponding undifferentiated cells do not show TREM2 genotype-dependent metabolic deficits. These energy deficits are not due to lower mitochondrial numbers, but may be influenced by enhanced mitochondrial superoxide levels. Indeed, mitochondrial impairment, namely energy deficiencies and aberrant ROS signalling, have been previously linked to a number of neurodegenerative diseases (Lin & Beal, 2006), most notably Parkinson’s disease (Park *et al*, 2018) but also AD and ageing (Trifunovic & Larsson, 2008; Sahin & Depinho, 2010; Wang *et al*, 2014). Our current findings are the first to show these metabolic deficits in disease relevant TREM2 hypomorphic microglia, however, whilst microglia upon activation can produce a burst of reactive oxygen species via the activity of the NAPDH oxidase (Bal-Price *et al*, 2002; Brown & Vilalta, 2015) it remains to be seen whether the production of cellular superoxide is altered in TREM2 hypomorphic microglia.

We identified aberrant responses following exposure to classical pro-inflammatory stimuli (LPS/IFNγ) in TREM2 hypomorphic iPS-Mg; specifically, reduced morphological changes and TNFα release. Both responses require rapid energy production which is produced when microglia undergo the metabolic switch to glycolysis (Finucane *et al*, 2019; Jiang *et al*, 2016). Glycolysis allows energy production and uptake of essential nutrients to support the rapid changes required by “activated” microglia in response to a stimulus, such as phagocytosis, proliferation, migration and induction of protein synthesis for cytokine and chemokine secretion (Vander Heiden *et al*, 2009; Shen *et al*, 2017; Gu *et al*, 2017; Culmsee *et al*, 2018). Indeed, we identified the ability of the TREM2 variant expressing iPS-Mg to undergo a normal switch in metabolism from a homeostatic, surveillance profile supported by oxidative phosphorylation, to one in which glycolysis is impaired. Interestingly, when we blocked glycolysis prior to LPS/IFNγ exposure, only control and R47H^het^ iPS-Mg were able to increase energy demand through enhanced oxidative phosphorylation. These data suggest a less dramatic loss of energy production in the AD-associated risk variant compared with the more severe NHD mutations.

The identification of comparable aerobic responses after glycolytic inhibition, but underlying metabolic and phenotypic differences between the control and patient AD R47H^het^ iPS-Mg, led us to probe the effect of glycolytic inhibition on cell stress pathways. Indeed, previous studies have shown that glycolytic inhibition can enhance oxidative stress (Shutt *et al*, 2010; Sharma *et al*, 2010), and with increased mitochondrial ROS levels in the TREM2 hypomorphic lines, we hypothesised that the discrepancies identified in the R47H^het^ lines may be due to a hyper-sensitivity to glycolytic inhibition. We found increased expression in the number of proteins indicative of metabolic and mitochondrial stress pathways in the R47H^het^ iPS-Mg when compared with control iPS-Mg supporting this hypothesis. Furthermore, when we investigated PGC1α, a master regulator of mitochondrial biogenesis and energy metabolism and signalling molecule downstream of a number of the pathways identified, we found enhanced negative regulation in the TREM2 hypomorphic lines. Collectively, these data suggest that whilst cell stress pathways are enhanced in R47Hhet lines during glycolytic inhibition, the cells’ ability to respond is downregulated. Indeed, when we looked at the transcriptional activity of PPARγ, a nuclear receptor and transcription factor co-activated by PCG-1α, we found that control iPS-Mg were able to respond to glycolytic inhibition whereas R47H^het^ and T66M^hom^ iPS-Mg were locked and unable to respond. PPARγ is widely expressed in inflammatory cells such as microglia and macrophages as well as adipose tissue where it controls inflammation, lipid metabolism and glucose homeostasis (Corona & Duchen, 2016). In support of the aberrant PPARγ signalling, we found protein levels were highly regulated by glycolytic inhibition, specifically in R47H lines and the heterozygous variant of T66M. Interestingly basal levels of PPARγ were significantly reduced in the severe NHD lines.

PPARγ, also known as the glitazone receptor or nuclear receptor subfamily 3 (NR1C3) is involved in insulin sensitisation and enhanced glucose metabolism, and plays a key role in cellular homeostasis (Tyagi *et al*, 2011). The prominent isoform expressed in inflammatory cells is PPARγ3. Pioglitazone, an agonist of PPARγ, has been shown to halt progression of Parkinsonism in a rodent model, ostensibly by inhibiting microglial inflammation and proliferation (Machado *et al*, 2019), although it is difficult to conclude if this is a direct effect on microglia given that the compound can affect a number of cells in these models. Targeting PPARγ for therapeutic benefit in AD has also recently been proposed (Khan *et al*, 2019). Interestingly we found that whilst the deficits in maximal respiration, glycolysis and glycolytic capacity observed in the R47H variants can be rescued by the PPARγ agonist, these deficits in the NHD hypomorphs cannot, suggesting that the reduced levels of PPARγ protein observed in these variants and the limited levels of mature TREM2 may have an uncoupling effect on the signalling to and from PPARγ. Furthermore, pre-incubating R47H^het^ iPS-Mg with pioglitazone prior to LPS/IFNγ exposure rescued the energetic metabolic phenotype, strongly linking PPARγ function to the locked immunometabolic switch observed in TREM2 hypomorphic lines.

PPARγ signalling has been identified as vital in cellular immune responses. Specifically, PPARγ activation has been shown to inhibit the expression of inflammatory cytokines and promote anti-inflammatory phenotypes (Ji *et al*, 2001; Clark, 2002). Whilst PPARγ expression is reduced in NHD TREM2 hypomorphs, one might expect a correspondingly higher level of TNFα secretion if PPARγ is controlling inflammation, however the opposite occurs. Whilst it may be true for TNFα, despite the requirement for the metabolic switch to glycolysis upon microglial activation for energy to produce cytokines and chemokines, we found previously that the ability of iPS-Mg to produce a whole range of cytokines was not greatly impaired when a stimulus of LPS alone was used (Garcia-Reitboeck *et al*, 2018). Here we primed with IFNγ and LPS, and were able to show a deficit in the secretion of TNFα in TREM2 hypomorphs compared with controls. TNFα and IL-6 expression may be controlled by activating transcription factor 3 (ATF3) following TLR stimulation or NFĸB, and at least three classes of transcription factor and coregulators are thought to control the inflammatory secretome (Medzhitov & Horng, 2009). Of further interest would be to determine the effects of multiple exposures to stimulants likely to evoke cytokine release in control iPS-Mg, to determine whether there are deficiencies or enhancements in TREM2 variant iPS-Mg.

To understand how PPARγ signalling influences the metabolic phenotype of the TREM2 variants we interrogated PFKFB3 signalling, a key regulatory enzyme involved in glycolytic induction (Rodríguez-García *et al*, 2017; Finucane *et al*, 2019) and identified as a target of PPARγ (Guo *et al*, 2010). We found that pioglitazone exerted its effects via a p38MAPK/PFKFB3 signalling cascade and we show that activation of the PPARγ/p38MAPK cascade and PFKFB3 activity is sufficient to rescue the functional deficit in Aβ_1-42_ phagocytosis identified in the TREM2 hypomorphic iPS-Mg, a key hallmark associated with AD pathogenesis. Finally, it is worth commenting on previous studies that identify p38- MAPK activation by pioglitazone (Xing *et al*, 2008; Ji *et al*, 2010). However, these studies use the compound at concentration 15 to 30x higher than our treatment, which itself is 7x lower than the reported EC50 for PPARγ activity.

In conclusion, we find that the presence of a TREM2 mutation detrimentally influences metabolic signalling by influencing not only basal levels of oxidative phosphorylation but also the requirement of a metabolic switch to glycolysis. This inability to switch on glycolysis during a change in environment significantly impacts upon the surveillant properties of homeostatic microglia leading to sub-optimal responses in key microglial functions such as phagocytosis. Our data highlight these deficits in a human, genetically disease relevant microglial model and uncover a significant dependence on p38MAPK signalling during glycolysis in iPS-Mg generated from R47H genotypes. This polymorphism seems to render the microglia hyper-sensitive to cellular stress, however this susceptibility enhances the effectiveness of PPARγ activation in upregulating cellular metabolism, leading to an ability to attenuate deficits in microglial function associated with disease pathogenesis.

## Acknowledgements

T.M. Piers was supported by funding to J.M. Pocock and J. Hardy from the Innovative Medicines Initiative 2 Joint Undertaking under grant agreement No 115976. This Joint Undertaking receives support from the European Union’s Horizon 2020 research and innovation programme and EFPIA. K. Cosker was supported by Eisai, working within the Eisai:UCL Therapeutic Innovation Group (TIG). A. Mallach was supported by the Biotechnology and Biological Sciences Research Council [grant number BB/M009513/1]. A. Mallach was funded by a LiDo Consortium PhD studentship. We would like to thank the patients and their families for their participation in this research project; E. Lohmann, Istanbul University, Istanbul Faculty of Medicine, Department of Neurology, Behavioral Neurology and Movement Disorders Unit, Istanbul 34390, Turkey; M. Blurton-Jones School of Biological Sciences, University of California, Irvine, San Diego; Henry Holden, and Pablo Garcia Reitboeck, UCL Queen Square Institute of Neurology for help with cells.

## Author contributions

T. M. Piers, J.M. Pocock and J. Hardy initiated the concept, T.M. Piers, K. Cosker and J.M. Pocock designed the experiments and T.M. Piers, K. Cosker, G. Thomas Johnson and A. Mallach carried out the experiments, R. Guerreiro analysed the in-house generated iPSC lines, J.M. Pocock, T.M. Piers and K. Cosker wrote the paper.

